# *Pseudomonas aeruginosa* virulence and biofilm factors provide independent routes for chronic colonization during catheter-associated urinary tract infection

**DOI:** 10.64898/2026.11.12.766126

**Authors:** Jared A. Carter, Vincent T. Lee

## Abstract

Bacterial colonization on indwelling medical devices such as urinary catheters is a leading cause of healthcare-associated infections and complicates patient treatment. The presence of a catheter in the urinary tract provides a foreign surface for biofilm formation and persistence in the host. For prominent uropathogens including *Escherichia coli*, chaperone-usher pili (CUP) are major colonization determinants for both non-catheterized and catheterized urinary tracts. However, the requirements for colonization in catheter-specific uropathogens such as *Pseudomonas aeruginosa* is less clear. Pseudomonal biofilms are promoted by many surface molecules including adhesins, motility apparatus, and polysaccharides. However, mutants lacking individual factors still formed biofilm during CAUTI suggesting there is functional redundancy. To resolve this functional redundancy for *P. aeruginosa* biofilm during CAUTI, we assessed a strain with chromosomal deletions of genes encoding eight known adhesins (Δ8) including flagella, type IV pili, five CUP, and exopolysaccharide for in vitro and in vivo biofilm formation. The Δ8 mutant showed the expected reduction in biofilm in vitro in laboratory media. However, this in vitro defect was reduced when a physiological medium such as urine was used. During CAUTI, Δ8 retained the ability to colonize in similar numbers on the catheter in mice. To determine if virulence and adhesins can function independently to promote biofilm during CAUTI, a strain that lacked both adhesins (Δ8) and type III secretion system (Δ8ΔT3SS) was tested. This double mutant showed >70-fold reduction for catheter colonization. We propose that *P. aeruginosa* employs multiple different strategies to form biofilms on medical devices to cause infections.

## INTRODUCTION

Healthcare-associated infections are largely caused by opportunistic pathogens that colonize the surfaces of an installed medical device ^1^. These infections are increasingly difficult to treat due to their ability to form biofilms. Catheter-associated urinary tract infections (CAUTI) are one of the most common healthcare-associated infections ^2–5^. CAUTI are a type of urinary tract infection that are complicated by the presence of the catheter. Without a catheter, the majority of urinary tract infections are caused by uropathogenic *E. coli* which require chaperone-usher pili (CUP) to attach to bladder epithelium ^6–8^. CUP binding promotes internalization into the host cell and the formation of biofilm-like intracellular bacterial communities that enhance evasion of host immunity ^9,10^. Uropathogenic *E. coli* remains the most prevalent causative agent of CAUTI and CUP remain important colonization determinants ^11,12^. The presence of a catheter provides an alternative reservoir for other microbes and increases the diversity of pathogens that cause CAUTI including *Pseudomonas aeruginosa* ^13^. In people without a catheter, *P. aeruginosa* causes less than 1% of urinary tract infections ^13^, but in people with catheter, *P. aeruginosa* causes over 10% of CAUTI ^14^. These clinical observations suggest that organisms that do not normally cause urinary tract infections are able to colonize the catheter to evade elimination by the host. This is further supported by imaging of catheters removed from CAUTI patients which have revealed bacterial biofilms to be abundant on the luminal surface of the catheter as well as the exterior surface proximal to the bladder ^15,16^. Thus, biofilm formation on the catheter surface is a key process for *P. aeruginosa* and other pathogens during CAUTI.

To understand how bacteria cause CAUTI, murine infection models have been established in which an implant is introduced at the same time as bacterial inoculation ^17^. Catheter insertion alone results in altered bladder histology and expression of pro-inflammatory cytokines ^18^. Subsequent infection with *P. aeruginosa* leads to weight loss that recover the weight after one week ^19^. These observations in animals are similar to clinical reports of patients infected by *P. aeruginosa* experience acute symptomatic that reduces in severity over time ^20^. After 14 days of infection, catheters removed from mice have been imaged by scanning electron microscopy and revealed that *P. aeruginosa* forms biofilm in the luminal surface of the catheter but was not visible on the external surface of the catheter ^21^. These results indicate that the murine model reproduces several aspects of clinical CAUTI. Studies using this CAUTI model have shown that *P. aeruginosa* requires the type 3 secretion system (T3SS) to cause acute virulence associated with weight loss, morbidity, and mortality ^19^. These mutants, though, retained the ability to form biofilm on the catheter, suggesting that additional factors contribute to biofilm formation.

*P. aeruginosa* biofilm formation has been extensively studied in vitro, typically in rich laboratory media using abiotic surfaces such as glass or plastic. As a species, *P. aeruginosa* encodes several extracellular factors that contribute to biofilm formation under these conditions including type IV pili, flagella, exopolysaccharides, and chaperone/usher pili (CUP). The model for biofilm formation ^22,23^ requires initial surface attachment, which is facilitated by type IV pili ^24^ and flagella ^25,26^. Subsequent increase in biofilm biomass utilizes exopolysaccharides, such as Psl and Pel ^27–29^. In addition to these factors, *P. aeruginosa* strains can encode five CUP (*cupA-cupE*) that contribute to in vitro biofilm ^30^ ^31–34^. Several of these mutants have been assessed in the murine CAUTI model, including mutants individually lacking flagella, type IV pili ^35^, quorum sensing ^36^, or exopolysaccharides ^21^. These studies show that each of these mutants were capable of colonizing the catheter and thus these factors are individually dispensable for *P. aeruginosa* CAUTI.

Because of the possible redundant functions of these biofilm factors, how *P. aeruginosa* forms biofilm during CAUTI remains unknown. To address this, we generated a mutant lacking flagella, type IV pili, CUP, and exopolysaccharides (Δ8) and tested the ability of this strain to form biofilm in vitro and colonize in a murine model of CAUTI. In vitro, this octuple mutant showed over 30-fold decrease in biofilm when grown in laboratory media as expected. When grown in more physiological relevant medium for infection, such as mouse or human urine, the biofilm defect was reduced to less than 3-fold. In vivo, we found that the Δ8 mutant was cleared more frequently by hosts than the wild-type strain, but it showed no difference in bacterial burden on the catheters of hosts that remained infected. In contrast, a Δ8ΔT3SS mutant which lacks both adhesins and T3SS activity showed a near 100-fold reduction in bacterial burden on the catheter of infected hosts. These findings suggest that catheter colonization in vivo can be facilitated by independent pathways that involve either bacterial adhesins or factors derived from T3SS-mediated host cell death.

## RESULTS

### Generation and confirmation of PA14 Δ8 mutant

We generated a PA14 Δ8 mutant lacking flagella, type IV pili, CUP, and exopolysaccharides and assessed phenotypes including growth in liquid culture (**Fig. S1**), swimming motility in low percentage agar (**Fig. S2**), twitch motility (**Fig. S3**), pyocyanin (**Fig. S4A**), and protease (**Fig. S4B**). Δ8 had similar growth in LB, pyocyanin production and protease secretion as wild-type PA14 (**Fig. S1**, **S2**, **S4A**, and **S4B**). The Δ8 mutant also had the expected defect in swimming for strains lacking *fliC* (**Fig. S2**) and defect in twitching for strains lacking *pilA* (**Fig. S3**). Further investigation revealed that the defect in *fliC* was complemented by trans expression of *fliC* from a plasmid (**Fig. S2**), while the *pilA* defect was partially complemented by trans expression of *pilA* from a plasmid (**Fig. S3**). These strains were further confirmed by genome sequencing, which verified the expected introduced gene deletions. Together, these data show that the Δ8 strain was generated with the intended genotype and corresponding phenotype.

**Figure 1.**
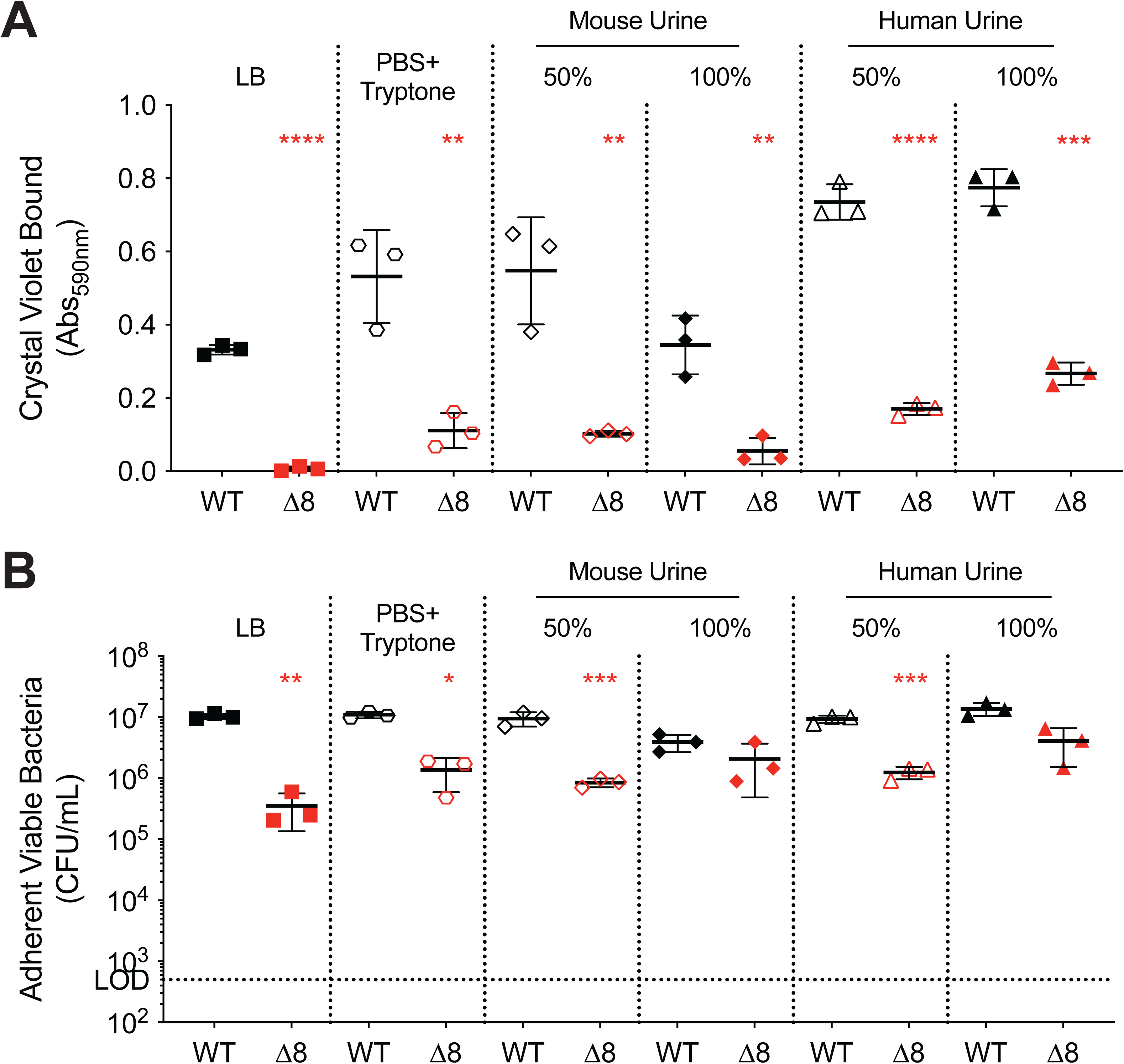
PA14 biofilm formation in urine differs from rich media. Biofilm formation on catheter material was performed by inoculating the lumen of 5 cm PE50 catheter tubing with 10^9^ CFU/mL of PA14 WT or Δ8 resuspended in the indicated media and incubated for 20hr at 37 °C in a humid chamber. After incubation, the lumen was eluted and washed with PBS followed by crystal violet staining of the catheter lumen (**A**) or detachment of bacteria from the catheter by homogenizing in PBS Tryptone (**B**). For crystal violet staining, reported is the Abs590nm normalized by subtracting staining of catheters with media alone. Symbols and bars represent the Average ± SD (**A**) or Geometric Average ± Geometric SD (**B**). Statistical significance was calculated in Prism 11 using Student’s t test for (**A**) or Lognormal Welch’s t test for (**B**). * P < 0.05, ** P < 0.01, *** P < 0.001, **** P < 0.0001

**Figure 2.**
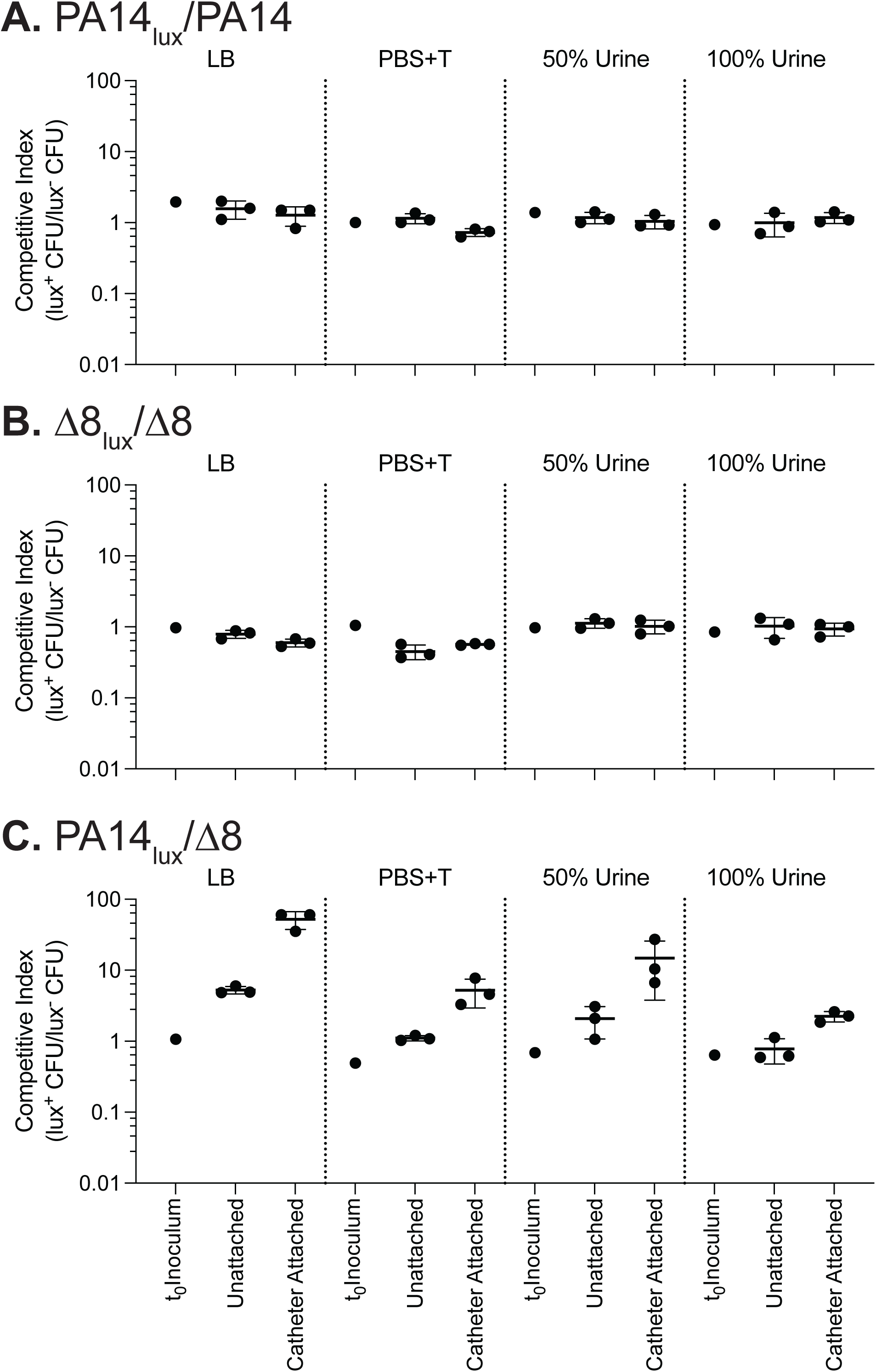
Δ8 has reduced attachment to catheter. Competitions assays were performed by mixing unlabeled PA14 and PA14_lux_ strains as **A** PA14_lux_/PA14, **B** Δ8_lux_/Δ8, or **C** PA14_lux_/Δ8 and resuspending the mixtures in LB, PBS+T, 50% mouse urine 50% PBS+T, or 100% mouse urine. Strains were mixed in a 1:1 ratio by volume, then inoculated in catheters and incubated at 37 °C for 20 hr in a humid chamber. CFU were determined in the inoculum at time of inoculation and bacteria in the flowthrough of the wash to determine ratio of non-adherent bacteria. Adherent bacteria were detached by homogenization. Plotted are the competitive indices determined as luminescent CFU compared to non-luminescent CFU with mean and standard deviation.

### Adhesins that contribute to PA14 biofilm are media dependent

Previous studies showed that *P. aeruginosa* forms biofilms in the catheter luminal surface but not the exterior surface during CAUTI ^21^. To mimic the in vivo condition, *P. aeruginosa* was resuspended in LB media, inoculated in the lumen of catheters used in CAUTI, incubated for 20 hours at 37 °C, then washed with sterile phosphate-buffered saline prior to the crystal violet assay ^37^. Wild-type PA14 formed detectable biofilms in the catheters (**Fig. 1A**), which agrees with previous studies showing biofilm formation on borosilicate glass, polystyrene, and polyvinyl chloride ^27^. In contrast, the Δ8 mutant formed low levels of biofilm, corresponding to >48-fold reduction in crystal violet staining compared to PA14 in LB (**Fig. 1A**). Crystal violet staining does not distinguish live and dead cells, so viable bacteria were assessed in similarly prepared biofilms. Bacteria were detached from catheters using a tissue homogenizer, then serially diluted and plated to enumerate viable bacteria as colony-forming units (CFU). In LB media, Δ8 showed 29-fold fewer CFU compared to PA14 (**Fig. 1B**). This suggests that results from the crystal violet assay are similar to CFU assay (within 2-fold). This reduction indicates that the Δ8 strain is more easily removed from the catheter by the wash, but nonetheless a subset of cells are still able to associate with the catheter.

In murine CAUTI models, strains are resuspended in phosphate-buffered saline and inoculated into the host ^19^. After inoculation, the host voids the inoculum and the remaining bacteria would be exposed to host urine. To understand how changes in media contribute to biofilm phenotype, PA14 and Δ8 were tested for biofilm formation in catheters in phosphate-buffered saline media supplemented with 1% tryptone (PBS+T). In PBS+T, Δ8 formed detectable biofilm that bound 4.8-fold less crystal violet than PA14 (**Fig. 1A**), and Δ8 showed 8-fold fewer viable bacteria than PA14 (**Fig. 1B**). We also tested biofilm formation in mouse urine diluted to 50% with PBS+T or in 100% mouse urine. In 50% mouse urine, Δ8 formed 4.9-fold less biofilm compared to PA14 and 11-fold reduction in viable bacteria, which was similar to the results in PBS+T. In 100% mouse urine, Δ8 showed 6.2-fold reduced biofilm by crystal violet assay, but only 1.9-fold reduction in viable bacteria. With increasing amounts of mouse urine in the media, the difference in biofilm formation between PA14 and Δ8 was reduced.

To determine if these phenotypes could be relevant in human infections, biofilm formation was tested in sterile-filtered pooled human urine. In human urine diluted to 50% with PBS+T, Δ8 showed 4.3-fold lower biofilm in crystal violet assays (**Fig. 1A**) and 7.5-fold reduction in viable bacteria compared to PA14 (**Fig. 1B**), similar to PBS+T. In 100% human urine, Δ8 showed a 2.9-fold reduction in crystal violet assay and 3.4-fold reduction in viable bacteria compared to PA14. In 100% mammalian urine, the difference in CFU between Δ8 mutant and PA14 was reduced, suggesting that the contribution of these known biofilm factors is media-dependent.

### PA14 does not outcompete Δ8 to form biofilm on catheters in urine

The small differences observed in biofilm formation between Δ8 and PA14 might be attributed to the bacteria being inoculated separately in different catheters. To determine whether PA14 can outcompete and exclude Δ8 mutant from the biofilm, competition experiments were performed by inoculating catheters with 1:1 mix of unlabeled strains with strains labeled with chromosomally encoded *lux* operon ^38,39^. The competitive index (CI) of the inoculum was assessed as a ratio of luminescent colonies divided by non-luminescent colonies. As a control, the CI at t_0_ for all combinations of luminescent and non-luminescent strains was less than 2-fold (**Fig. 2A-C** and **Fig. S5**). After incubation for 20 hrs, the CI was determined for two populations: (i) unattached bacteria in the lumen (CI_lumen_) and (ii) bacteria attached to the surface of the catheter (CI_catheter_). The unattached bacteria in the CI_lumen_ indicated differences in growth between the strains. For the PA14_lux_ vs PA14 mixture in LB, there was no competition as expected (CI_lumen_ was 1.6 and CI_catheter_ was 1.3) (**Fig. 2A**). Similar results were observed when strains were tested in PBS+T, 50% mouse urine, and 100% mouse urine (CI_lumen_ ranged 1.0 to 1.2 and CI_catheter_ ranged 0.8 to 1.3). The Δ8_lux_ vs Δ8 mixture showed similar results with CI of ∼1 across all media and sample locations (**Fig. 2B**). These results demonstrate that strains labeled with chromosomal *lux* did not introduce a growth difference to *P. aeruginosa*.

For the PA14_lux_ vs Δ8 mixture in LB, PA14_lux_ slightly outcompeted Δ8 in the lumen (CI_lumen_ = 5). In contrast, PA14_lux_ outcompeted Δ8 in the biofilm on the catheter (CI_catheter_ = 52). When tested in different media that reflect CAUTI, the advantage of PA14 over Δ8 in the lumen was reduced in PBS+T, 50% urine, and 100% urine (**Fig. 2C**). The difference between PA14 and Δ8 in the biofilm on the catheter was also reduced (**Fig. 2C**). When the *lux* operon was placed on Δ8 and the experiments repeat with mixtures of Δ8_lux_ and PA14, complementary results were obtained (**Fig. S5**). Additionally, similar findings were observed in assays conducted using human urine (**Fig. S6A-B**). Together, these results support that the known adhesins contribute to attachment by PA14 in laboratory media which differs from Δ8. However, in urine media, the difference between PA14 and Δ8 was reduced likely due to other factors mediating attachment.

### Adhesins contribute to colonization during chronic CAUTI

To understand if the 8 adhesins contribute to catheter colonization in vivo, Δ8 was tested in a murine model of CAUTI. Mice were transurethrally catheterized and infected with 3×10^7^ CFU of Δ8 and monitored for symptoms and weight loss as previously ^19^. Compared to the previously published PA14 data, Δ8 showed no statistical difference in the first three days post-infection in terms of mortality (**Fig. 3A**). From day 3-14, Δ8 caused a modest reduction in mortality that is not statistically significant (P = 0.1352 by Mantel-Cox) (**Fig. 3A**). For average weight of mice that survived over the 14-day experiment, mice infected with PA14 appeared to show more weight-loss compared to mice infected with Δ8, but this difference was only statistically significant on Days 4 and 7 (**Fig. 3B**). We tested whether Δ8 mutant acquired a loss in cytotoxicity against mammalian cells (**Fig. S7**). However, Δ8 showed reduced cytotoxicity toward macrophages and urothelial cells associated with the loss of both flagella and type IV pili (**Fig. S7A** and **S7B**). These results suggest that the 8 adhesins may have a minor contribution to acute virulence.

**Figure 3.**
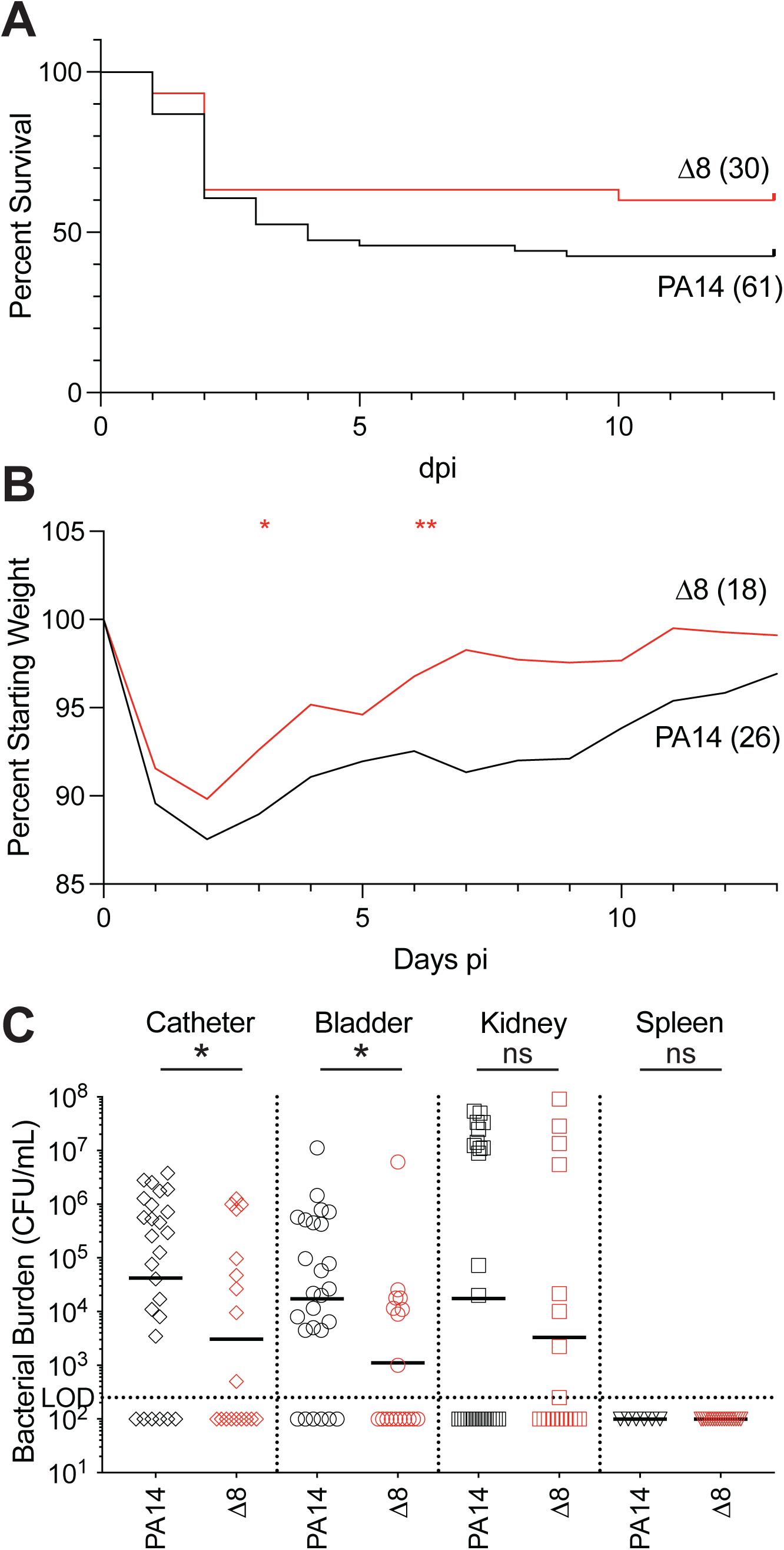
Biofilm adhesins have minor contribution to virulence. Mice were transurethrally catheterized and infected with 3×10^7^ CFU of indicated strain. Over 13 -15 days post-infection, mice were assessed for signs of acute infection including lethargy, ruffled fur, and weight-loss exceeding 20%; mice showing these signs were euthanized and reported in (**A**). Statistical significance was assessed using Mantel-Cox test in GraphPad Prism 11; the comparison was not significant (P > 0.05). Daily weight was measured until euthanization or infection endpoint (13-15 dpi); relative weight-loss of those that survived to infection endpoint are reported in (**B**). Statistical significance was determined using two-way ANOVA with Šídák test for multiple comparison. At 14 dpi, geometric average CFU was determined in the indicated sites (**C**). Statistical significance was determined using Mann Whitney U test * P < 0.05, ** P < 0.01

To assess whether the Δ8 mutant’s slight reduction in virulence is due to reduced colonization in the host, we assessed CFU at chronic timepoints where mice survived to the infection endpoint (13-15 dpi). Δ8 showed a statistically significant 1-log reduction in average CFU in the catheter and bladder compared to the previously published PA14 data (**Fig. 3C**). Δ8 showed a 5-fold reduction in average CFU in the kidneys, which was not statistically significant (P = 0.5550). Previous results showed no *P. aeruginosa* in the spleen in animals infected with PA14 after 1 week of infection ^19^. Δ8 showed similar absence of bacteria in the spleen (**Fig. 3C**). These results suggest that one or some combination of these 8 adhesins contributes persistence of *P. aeruginosa* PA14 in bladder and catheter. As Δ8 caused chronic colonization of the catheter in some hosts, there are likely additional factors contributing to its persistence.

### Virulence contributes to Δ8 colonization during chronic CAUTI

Our previous work showed that acute virulence (morbidity, mortality, and weight loss), is driven primarily by T3SS, whereas chronic colonization of the catheter and bladder is independent of T3SS. To determine if T3SS and the adhesins work synergistically to contribute to Δ8 colonization, we generated a Δ8 that lacks a functional T3SS (Δ8Δ*pscC*) to determine if it would further reduce colonization of the urinary tract by eliminating contribution of acute virulence. PA14 Δ8Δ*pscC* strain has reduced virulence with a low mortality phenotype and weight loss profile that was similar to previous data from Δ*pscC*Δ*xcp* (**Fig. 4A and 4B**) ^19^. For chronic colonization assessed at 2 weeks post infection, the Δ8Δ*pscC* showed a 1-log reduction in average CFU in the catheter and bladder (**Fig. 4C**).

**Figure 4.**
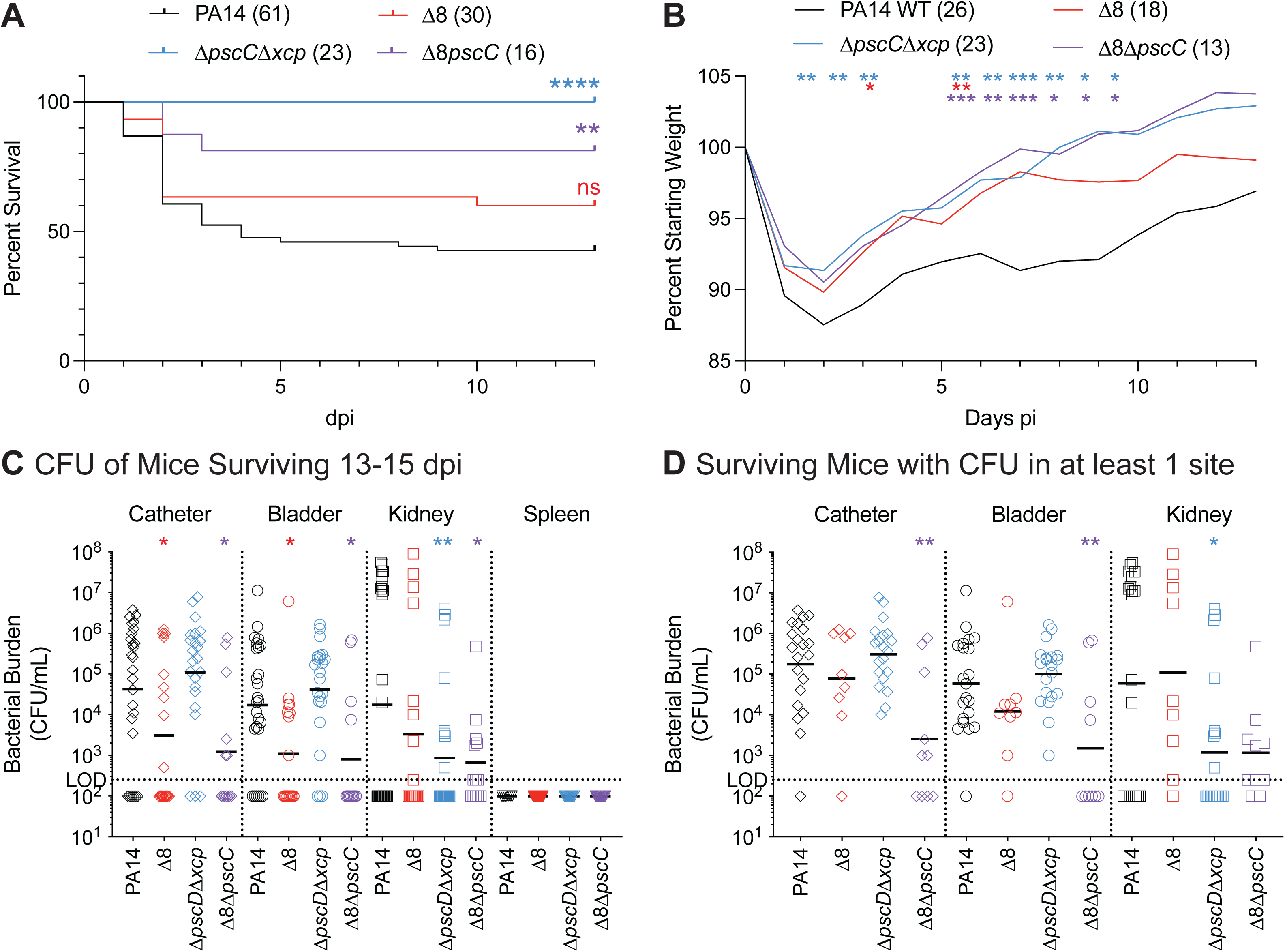
Adhesins and T3SS provide independent pathways for chronic catheter colonization. Mice infected with indicated strains were monitored over 13 -15 days post-infection, and mice were assessed for signs of acute infection including lethargy, ruffled fur, and weight-loss exceeding 20%; mice showing these signs were euthanized and reported in (**A**). Statistical significance was assessed using Mantel-Cox test in GraphPad Prism 11. Daily weight was measured until euthanization or infection endpoint (13-15 dpi); relative weight-loss of those that survived to infection endpoint are reported in (**B**). Statistical significance was determined using two-way ANOVA model with Šídák test for multiple comparison. At 14 dpi, mice were sacrificed and geometric average CFU was determined in indicated site (**C**). Geometric average CFU from mice that had detectable CFU in at least one site are shown for reanalysis (**D**). Statistical significance was determined using Kruskal-Wallis test with comparisons made to PA14; * P < 0.05, ** P < 0.01, *** P < 0.001, **** P < 0.0001.

Further inspection of the data show that there is a bimodal distribution of bacterial burden of *P. aeruginosa* on the catheter among different animals. This heterogeneity could be explained by the varied host response to infection expected from genetically heterogenous outbred mice. From our observations, while all animals respond to infection by voiding the bladder to eliminate the pathogen, there is large variation between animals in the timing and volume of voiding. This is supported by statistical analysis that geometric standard deviation factor for PA14 is 47 which is quite large. Furthermore, the P values comparing the average CFU of PA14 WT to Δ8 and Δ8Δ*pscC* in the catheter are close to 0.05 (0.04 and 0.01, respectively) which is often considered significant. Thus, the analysis that combine the results of animals that cleared infection with those that remain infected confound our analysis.

To deconvolute the analysis of these data, we reanalyzed the infection data by removing data for animals that fully clear the infection as defined as having no detectable *P. aeruginosa* in any of the sampled tissues (**Fig. 4D**). For PA14 WT and Δ*pscC*Δ*xcp*, more than 80% of animals remained infected (21/26 and 20/23, respectively). For Δ8, on 50% of animals remained infected (9/18). Surprisingly, 11/13 animals remained infected for Δ8Δ*pscC*. Analysis of animals that remain infected reduced the heterogeneity as the geometric standard deviation factors reduced from 47 to 15 for wildtype PA14 and from 53 to 27 for Δ8. For animals that remain infected at the 2-week experimental endpoint, Δ8 only shows a 2-fold reduced average CFU in the catheter (**Fig. 4D**). When a similar analysis was performed for Δ8Δ*pscC*, mice that remain infected after 14 days showed a 71-fold reduction in average CFU on the catheter compared to WT-infected mice. These results indicate that *P. aeruginosa* can employ either extracellular biofilm factors or cytotoxic virulence factors to form biofilm during CAUTI. Importantly, the absence of both extracellular biofilm factor and T3SS allows the host to effectively eliminate the majority of bacteria from the urinary tract.

### Flagella, pili, and polysaccharides all contribute to PA14 colonization in chronic CAUTI

We next sought to determine which of the eight extracellular biofilm factors are important for chronic colonization during CAUTI. Since acute virulence is dispensable for chronic colonization, the Δ*pscC* background was selected to generate mutants lacking either all 5 CUP (Δ5Δ*pscC*) or flagella, type IV pili, and exopolysaccharide (Δ3Δ*pscC*). At 14 dpi, the Δ5Δ*pscC* mutant showed a 4-fold and 8-fold reduction on the catheter and in the bladder, respectively (**Fig. S8A**). The Δ3Δ*pscC* mutant showed 11-fold and a 12-fold reduction in the catheter and bladder, respectively (**Fig. S8A**). While ≥80% of mice remained infected in one or more tissue, more mice cleared Δ3Δ*pscC* from the catheter than Δ5Δ*pscC* (4/16 vs 1/16). To understand whether there is a difference between the Δ5Δ*pscC* and Δ3Δ*pscC* mutants, a similar analysis was performed as in **Fig. 4D** to remove mice that fully cleared infection and focus on difference in bacterial burden of mice that did not clear infection. When compared to WT, Δ5Δ*pscC* mutant showed a 5-fold and a 12-fold reduction on the catheter and in the bladder, respectively (**Fig. S8B**). The Δ3Δ*pscC* mutant showed a 22-fold reduction and 26-fold reduction on the catheter and in the bladder, respectively (**Fig. S8B**). Though neither mutant fully phenocopies Δ8Δ*pscC*, Δ3Δ*pscC* has a more pronounced defect than Δ5Δ*pscC*. These results suggest that all of the extracellular biofilm factors contribute to colonization in the CAUTI model.

## DISCUSSION

Previous works in the murine CAUTI model have assessed the contribution of extracellular biofilm factors individually including exopolysaccharides ^21^, quorum sensing ^36^, flagella and type IV pili ^35^. Mutants lacking each of these individual factors retain their ability to colonize the catheter in vivo, but this may be due to functional redundancy in which the absence of any individual factor is masked by the remaining surface molecules. To address this possibility, we took the approach of generating a PA14 mutant that lacks 8 extracellular biofilm factors that contribute to in vitro biofilm formation.

The Δ8 mutant was tested in vitro biofilm grown in LB using the crystal violet binding assay and the results confirmed the expected biofilm phenotype for PA14 Δpel previously described for other abiotic surfaces ^28^. However, when tested in more physiologically relevant media to CAUTI (such as urine), the large difference in bacterial adherence between PA14 and Δ8 observed in LB became less than 5-fold. In competition assays, Δ8 was outcompeted by wildtype in LB but was able to colonize within 5-fold when grown in urine (**Fig. 2C**). These findings indicate that biofilm formation in urine can require additional bacterial factors in addition to the eight known factors. In testing the contribution of the 8 adhesins in the murine CAUTI model, we found that Δ8 shows a 10-fold reduction in average CFU on the catheter compared to wild-type PA14 (**Fig. 3C**). We also found that Δ8 was cleared more frequently from infected mice than wild-type PA14. Upon reanalysis after removing data from animals that cleared *P. aeruginosa*, the bacterial burden of Δ8 on the catheter in mice was similar to the wild-type PA14 (**Fig. 4D**). The colonization of the catheter during CAUTI by either a T3SS mutant ^19^ or Δ8 raised an intriguing possibility that these two processes acted independently to promote biofilm formation. To test this possibility, we generated the Δ8Δ*pscC* mutant and verified that it had reduced acute virulence during CAUTI (**Fig. 4A-B**). Most interestingly, the Δ8Δ*pscC* mutant had a near 2-log decrease in average CFU for mice that did not clear the infection (**Fig. 4D**). These results demonstrate the ability of *P. aeruginosa* to employ either cytotoxic virulence factors or extracellular biofilm factors to promote colonization in a device-associated infection.

How do these data inform the complicated infections of the urinary tract? There are numerous dynamic factors during CAUTI that may influence biofilm formation on the catheter including timing and volume of bladder voiding, the host immune response, the bacterial virulence factors that counteract the host immune response, and the interactions of each of the factors with the catheter surface. We propose a model in which *P. aeruginosa* enters the bladder and the host responds with infiltration of immune cells (**Fig. 5A**) ^20^. In the absence of a catheter, *P. aeruginosa* are eliminated by the host immunity and/or voiding of the bladder unless they can ascend to the kidneys to cause pyelonephritis ^20^. In the presence of catheters, *P. aeruginosa* can attach to the luminal surface and can resist host immune response and elimination due to bladder voiding. *P. aeruginosa* mutants lacking T3SS colonize equally well on the catheter after 2 weeks as wild-type PA14 ^19^, indicating that the adhesins alone can mediate attachment to the catheter and form biofilms that can counter the host immune response. In this study, mutants lacking the surface adhesins (Δ8) were still able to colonize the catheter in similar numbers, albeit the bacteria were cleared more often than wild-type PA14. This suggests T3SS provides a second path for catheter colonization that is independent of surface adhesins. PA14 T3SS causes host cell death associated with ExoU activity. Cell death can release host components that coat the catheter surface and provide additional molecules for *P. aeruginosa* to adhere and grow as a biofilm (**Fig. 5B**). Once they form a biofilm, Δ8 can continue to deliver type III effectors to counter host immunity. For Δ8 ΔT3SS which lacks both pathways, the bacterial numbers on the catheter were reduced by nearly 2-logs supporting the presence of two independent pathways for chronic CAUTI.

**Figure 5.**
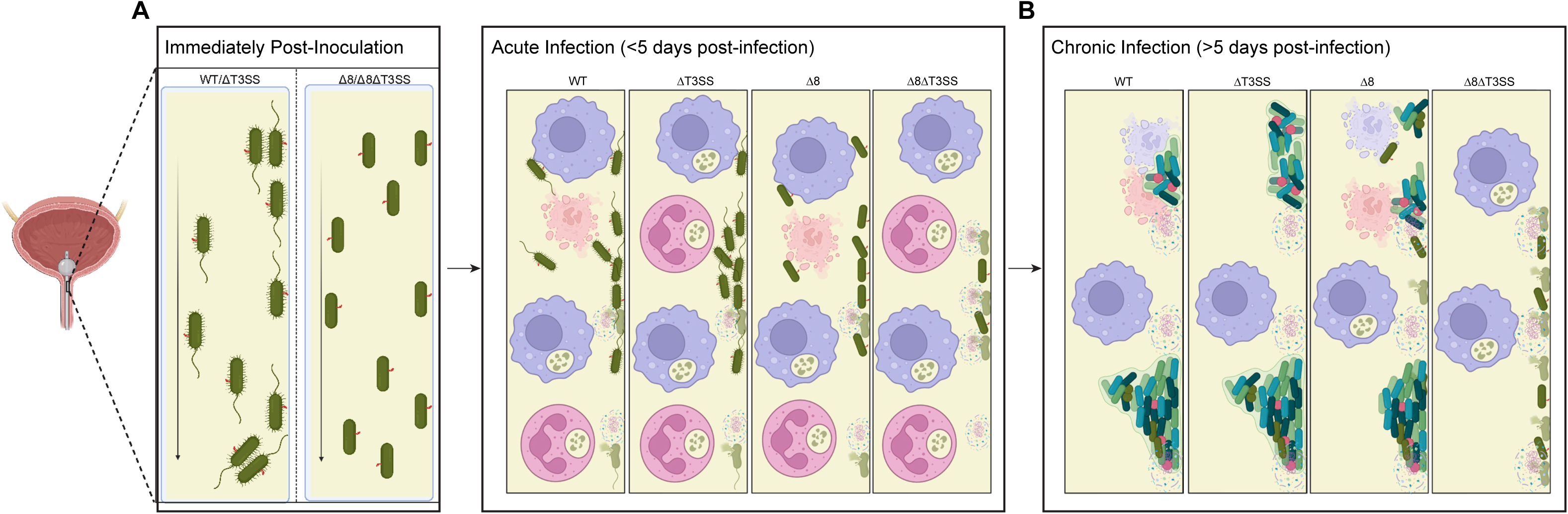
Model of *P. aeruginosa* colonization during CAUTI.

These findings raise the question of whether other CAUTI pathogens also require both biofilm adhesins and cytotoxic factors during chronic biofilm infections. The ability to bind host proteins is a shared feature among common CAUTI pathogens including uropathogenic *E. coli*, *Enterococcus*, *Staphylococcus, Klebsiella*, and *Acinetobacter* ^18,40,41^. As the preeminent CAUTI pathogen, uropathogenic *E. coli* requires CUP in CAUTI in order to bind host tissue as well as the catheter during infection ^11^. *Enterococcus faecalis* encodes Ebp pili to promote biofilm formation and fibrinogen binding ^42^. Staphylococci encode numerous surface proteins that bind host matrix molecules including fibrinogen-binding proteins, which contribute to catheter colonization during a muroid heart valve model ^43,44^. *Klebsiella* encode type 1 and type 3 pili which independently contribute to urinary tract colonization in the presence or absence of a catheter ^45,46^. *Acinetobacter* encodes two CUP, which both contribute to catheter colonization in vivo ^47^. From studies of other CAUTI pathogens, we note that a common factor that promotes in vivo biofilm formation is the ability to bind released host components, such as fibrinogen, which deposit on the catheter ^41^. However, the magnitude of contribution for mutants lacking specific adhesins varies between organisms. One possibility is that some organisms encode additional factors that induce a more inflammatory environment, thereby increasing the diversity of host proteins released from lysed cells that promote bacterial colonization. Bacterial factors that lyse host cells have been identified in strains of *Enterococcus* ^48^ ^49^, *Staphylococcus* ^50^, *Klebsiella* ^51^, and *Acinetobacter* ^52^, but further research is needed to understand how these virulence and adhesive factors synergistically can contribute to chronic colonization.

## METHODS

### Bacterial strains

*Pseudomonas aeruginosa* PA14 and strains with specific gene deletion used in this study are listed in **Supplemental Table 1**. In-frame deletion constructs of *fliC*, *pilA*, *pscC* genes as well as *pel*, *cupA*, *cupB*, *cupC*, *cupD*, and *cupE* operons were made using primers listed in **Supplemental Table 2**. The 1 kb regions upstream and downstream of the gene or operon were amplified by PCR and cloned into pCR-blunt (Invitrogen). After sequence verification, the two 1 kb fragments were subcloned into a pEX-Gn vector to generate a construct in which the gene or operon of interest is deleted ^53^. Each in-frame deletion was introduced into *P. aeruginosa* PA14 by conjugation using SM10 λpir harboring the pEX-Gn deletion plasmid. Cointegrants were selected on LB with 25 µg/mL Irgasan and 75 µg/mL gentamicin. Resolved deletion strains were counter-selected on LB agar with 10% sucrose. Strains were confirmed to regain gentamicin sensitivity and the absence of the genetic loci were confirmed by whole genome sequencing (SeqCenter, LLC). Genomic sequences of the strains generated in this study are archived in the NCBI SRA database under BioProject PRJNA1556129.

To generate *lux* strains, the miniCTX plasmid encoding the *lux* operon ^38,39^ was introduced into PA14 wild-type or Δ8 by conjugation ^53^. Cointegrants were selected on LB agar supplemented with 75 µg/mL tetracycline and 25 µg/mL Irgasan. Cointegrants were then conjugated with pFLP2 plasmid ^38^ and selected on LB agar supplemented with 150 µg/mL carbenicillin and 25 µg/mL Irgasan. Strains were then resolved by selection on LB agar with 10% sucrose. Resolved strains were confirmed to regain sensitivity to carbenicillin and tetracycline. Luminescence was confirmed using iBright 1500 with 90 second exposure time.

For swim motility complementation experiments, the in-frame PA14 *fliC* gene was cloned into pMMB and transformed into strains via conjugation. For twitch motility complementation experiments, the PA14 *pilA* gene including the 1 kb upstream intergenic region was cloned into a promoter-less pMMB plasmid and transformed into strains via conjugation.

For bacterial propagation prior to in vitro assays and infections, all strains were grown in LB media with aeration at 37 °C.

### Growth assay

In triplicate, overnight cultures of strains were diluted 1:100 in a final volume of 180 µL. Strains were incubated at 30°C with continuous shaking and absorbance at 600 nm was measured every 15 minutes over a 16 hr period. For plotting, at each timepoint the OD_600_ for bacterial culture was subtracted by OD_600_ of uninoculated media at the corresponding time.

### Agar Motility Assays

For swim motility assays, cultures were grown overnight in LB supplemented with 150 µg/mL carbenicillin. In triplicate, 0.5 µL of overnight culture was used to inoculate a 25 mL LB 0.3% agar plate supplemented with 50 µg/mL carbenicillin and 1 mM IPTG. Plates were incubated for 24 hr at room temperature, then colony diameter was measured using ImageJ.

For twitch motility assays, overnight cultures were stab inoculated on 5 mL LB 1% agar plates in triplicate. Plates were incubated for 24 hr at 30°C, then colony area was measured using ImageJ.

### Pyocyanin Assay

Strains were grown overnight in 5mL LB and OD_600_ was measured. 1.5mL of bacterial culture was centrifuged and the supernatant was treated with 0.5 mL of chloroform to isolate pyocyanin. Pyocyanin was extracted from the organic layer with addition of 0.25 mL of 0.2N HCl. 0.2 mL of extracted pyocyanin was used to measure absorbance at 520 nm. Levels of pyocyanin were normalized by bacterial culture density. Experiments were performed in triplicate.

### Protease Secretion Assay

Strains were grown to OD_600_ 0.2 and 2µL of culture were spotted on milk agar. Plates were incubated at 37°C for 18 hr, then the diameter of the optically clear region surrounding the inoculation point was measured using ImageJ.

### Biofilm assay

Single colonies of bacteria were resuspended in LB media and grown ∼6 hr at 37 °C until cultures reached OD_600_ = 1.0. Following growth, the equivalent of 10^9^ CFU were pelleted and washed once with 10 mM MOPS media. Cells were resuspended in the specified test media at a concentration of 10^9^ CFU/mL and injected into the lumen of UV-sterilized 5 cm polyethylene catheter tubing (0.023” inner diameter) via micropipette. Alternatively, catheters were filled with media alone to determine background effects. This inoculation delivers 15 µL of resuspension to the catheter, filling the catheter end-to-end. Catheters were incubated in a humid container at 37 °C for 20 hours.

Experiments to determine crystal violet binding and viable bacteria in **Fig. 1** were perfomred concurrently using the same bacterial inoculum for a given medium. To perform crystal violet assays ^37^, the lumen of the catheter was washed once using PBS+T then voided over a clean paper towel. Catheters were filled end-to-end with 0.1% Crystal Violet solution and stained for 10 minutes. Following staining, catheters were voided over a clean paper towel and washed two times with double-deionized water to remove excess dye. Bound crystal violet was dissolved in 20 µL of 30% acetic acid and eluted into a well of 96 well microtiter plate containing 180 µL of 30% acetic acid. The plate was incubated shaking at room temperature for 5 minutes then absorbance at 590 nm was measured. Values reported in the figure represent Absorbance subtracted by background staining caused by media alone.

For determination of viable bacteria following the 20 hr incubation, the lumen of the catheter was washed out using PBS+T and the catheter was aseptically cut and placed in 1 mL PBS+T to detach viable bacteria using a tissue homogenizer (“Catheter Attached” samples). Samples were serially diluted and plated on LB agar to determine CFU.

For competition assays, strains were luminescent labeled with the *lux* operon at the *ctx* site ^38^. After strains reached OD_600_ 1.0, the equivalent of 10^9^ CFU were washed and resuspended to 10^9^ CFU/mL as above, then the strain resuspensions were mixed in equal volume to make a single inoculum. The inoculum was serial diluted and plated on LB to enumerate total CFU. Lux^+^ colonies were identified by luminescence using an iBright 1500 with 90 second exposure time. Following the 20-hour incubation, the lumen of the catheter was washed out using PBS+T and the viable bacteria in this wash (“Unattached” samples) was determined. Then, the catheter was aseptically cut and placed in 1 mL PBS+T to detach viable bacteria using a tissue homogenizer (“Catheter Attached” samples). Samples were serially diluted and plated on LB agar to determine total and Lux^+^ CFU in the Unattached and Catheter Attached samples.

### LDH Release Assay against mammalian cell culture

J774 macrophages were propagated in Dulbecco’s Minimal Essential Media supplemented with 10% fetal bovine serum and phenol red. T24 urothelial cells were propagated in McCoy’s 5a Modified Medium supplemented with 10% fetal bovine serum and phenol red. Mammalian cell lines were incubated overnight at 37 °C, 5% CO_2_ and passaged no more than 10 times. The day before cytotoxicity assays, cells were trypsinized, counted, and seeded in a 96-well format at 4×10^4^ cells/well in 100 µL of fresh propagation media and incubated overnight at 37°C, 5% CO_2_. Prior to infection, cells were washed with phosphate-buffered saline two times, either cell type was resuspended in Dulbecco’s Minimal Essential Media without fetal bovine serum or phenol red, and cells were acclimated for 2 hr at 37°C, 5% CO_2_. Overnight cultures of indicated bacterial strains were subcultured 1:100 and grown to OD_600_ 0.2.

In triplicate, cells were infected using indicated bacterial strains at a MOI of 20 and plates were centrifuged at 4000 x g for 1 min to bring bacterial strains in proximity to mammalian cells. Plates were incubated for 3 hr at 37°C, 5% CO_2_. Following the incubation period, 50 µL of supernatant from the infection plates were transferred to a new plate, mixed 1:1 with Cytotoxicity Detection Kit (Roche), and developed for 10-15 minutes. The absorbance of the mixture was then measured at 492 nm. To calculate the % cytotoxicity, the 492 nm reading for no treatment was set to 0% cytotoxicity (Min) and treatment with 2% Triton X-100 was set 100% cytotoxicity (Max) using the equation below:

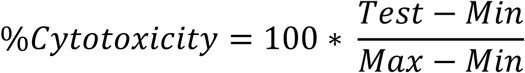

### Murine CAUTI Model

19-21 g CF-1 outbred female mice were obtained from Charles River, Wilmington, MA. Mice were housed in a group of five and acclimated for seven days. The weight of the mice on the day of infection is 23-29 g and infected in groups of 5 or 10 for each cohort. All procedures were performed in accordance with protocols approved by the IACUC at the University of Maryland at College Park (protocol R--MAY-24-15). The CAUTI model was used as previously described ^21^ ^19^. Mice were anesthetized with ketamine (144 mg/kg body weight) and xylazine (14.4 mg/kg body weight). A polyethylene catheter with size PE50 were cut into 5 cm guide or 0.4 cm implant (Braintree Scientific, Inc.). The catheter pieces were treated with ethyl acetate for 10 min to round edges and dried overnight to remove solvent. All catheters were sterilized under 254 nm UV light (UVP, 15 W) at 10 cm distance for 10 min. The periurethral areas were decontaminated with 70% isopropanol. A 0.4 cm implant was loaded onto a sterile 10 cm metal stylet followed by the 5 cm guide. After coating the stylet, implant and guide with lubricant, the metal stylet was placed in the urethral opening and inserted into the bladder. Deposition of the implant was achieved by advancing the 5 cm guide over the stylet and into the bladder. While the 5 cm guide remains in the bladder, the metal stylet was removed and the bladder was gently depressed to void the bladder of urine through the guide. The bacteria were grown to OD_600_ of 1.0, spun down and resuspended in PBS with 1% tryptone to a final concentration of 1×10^9^ CFU/mL. Bacteria were loaded into a 1 mL syringe with 23-gauge needle. The syringe is attached to the 5 cm guide catheter and 35 µl of the inoculum was injected into the bladder for a total inoculum of 3.5 x 10^7^ CFU. The tubing was subsequently removed. Mice were injected intraperitoneally with 100 µL of 100 mg/mL ampicillin eliminate non-*P. aeruginosa* introduced during the procedure. Mice were monitored daily for 14 days for weight and clinical symptoms. At the time of sacrifice or end of experiment, animals were dissected. The implant and indicated organs were dissected from each mouse, re-suspended in 1 ml sterile PBS-T, homogenized with tissue homogenizer (Omni International, USA), and serial dilutions were plated on LB agar for CFU enumeration. Animals lacking the implant were excluded from survival and CFU analyses. Animals that were deceased rather than sacrificed were dissected to verify the animal retained the catheter. The animals that retained the catheter were included in survival curve analysis, but excluded from CFU determination.

## ACKNOWLEDGEMENTS

We thank Sao-Mai Nguyen-Mao for generating PCR fragments for deletion of each of the adhesins and generation of individual mutants of cup. We thank funding from the NIH (T32 AI089621 for J.A.C. and R01 AI110740 for V.T.L.). We thank members of the lab for critical reading of the manuscript.

## Supplemental Figures

**Figure S1 Δ8 grows similar to PA14 wild-type in liquid media.** In triplicate, strains were grown in LB for 16 hr at 30°C. Absorbance at 600 nm was measured every 15 minutes. Symbols represent the triplicate average and bars represent SD.

**Figure S2 Δ8 lacks swim motility phenotype.** 0.5µL of overnight culture of indicated strains was inoculated on LB 0.3% agar supplemented with 150 µg/mL carbenicillin and 1 mM IPTG and incubated for 24 hr at room temperature. (**A**) Representative image of swim motility phenotypes. (**B**) Colony diameter was measured from triplicate experiments using ImageJ. Symbols represent the triplicate average and bars represent SD.

**Figure S3 Δ8 lacks twitch motility phenotype.** PA14 strains harbor either a promoter-less pMMB vector or pMMB pilA with native promoter. Overnight cultures of strains were stab inoculated on LB 1% agar supplemented with 150 µg/mL carbenicillin and incubated at 30°C for 24 hr. The colony area was measured from triplicate experiments using ImageJ. Symbols represent the triplicate average and bars represent SD.

**Figure S4 Expression of pyocyanin and protease by PA14 mutants. A** Indicated strains were grown overnight in LB and the supernatant pyocyanin was chloroform extracted. Extracts were treated with 0.2N HCl and the absorbance at 520 nm was measured and normalized by bacterial culture density at 600 nm. **B** Strains were grown to OD_600_ 0.2 and 2µL of culture was spotted on milk agar and incubated for 18 hr at 37°C. The area of optically cleared agar around the inoculation site was measured using ImageJ. Symbols represent the triplicate average and bars represent SD. Unpaired t tests with comparisons to WT were performed in Prism 11 to determine significance ** P < 0.01, *** P < 0.001

**Figure S5. Δ8 has reduced attachment to catheter.** Competitions assays were performed by mixing unlabeled PA14 and PA14 Δ8_lux_ strains in a 1:1 ratio by volume, then inoculated in catheters and incubated at 37 °C for 20 hr in a humid chamber. CFU were determined in the inoculum at time of inoculation and bacteria in the flowthrough of the wash to determine ratio of non-adherent bacteria. Adherent bacteria were detached by homogenization. Competitive index was determined as luminescent CFU compared to non-luminescent CFU. Symbols represent the triplicate average and bars represent SD.

**Figure S6. Δ8 has reduced attachment to catheter in human urine.** Competition assays were performed by resuspending PA14 or PA14_lux_ strains in (**A**) 50% Human Urine or (**B**) 100% Human Urine. Strains were mixed in a 1:1 ratio by volume, then inoculated in catheters and incubated at 37 °C for 20 hr in a humid chamber. CFU were determined in the inoculum at time of inoculation and bacteria in the flowthrough of the wash to determine ratio of non-adherent bacteria. Adherent bacteria were detached by homogenization. Competitive index was determined as luminescent CFU compared to non-luminescent CFU. Symbols represent the triplicate average and bars represent SD.

**Figure S7 Cytotoxicity against mammalian cells by PA14 mutants. A** J774 macrophages or **B** T24 bladder cells were infected with indicated midlog culture strains at a multiplicity of infection of 20 for 3 hr, then cytotoxicity was measured using lactate dehydrogenase release assay and normalized by 100% release by treatment with 2% Triton X-100. Symbols represent the triplicate average and bars represent SD. Unpaired t tests with comparisons to WT were performed in Prism 11 to determine significance * P < 0.05, ** P < 0.01, *** P < 0.001, **** P < 0.0001

**Figure S8. CUP, flagella, type IV pili, and pel contribute to colonization.** CUP, flagella, type IV pili, and pel contribute to colonization. Mice were infected with PA14 strains lacking 8 adhesins, all 5 CUP (Δ5), or flagella/type IV pili/pel (Δ3). At 14 dpi, mice were sacrificed and geometric average CFU was determined in indicated site (**A**); mice with detectable CFU in at least one site are presented further analysis (**B**).

**Supplemental Table 1. Bacterial strains used in this study**

**Supplemental Table 2. Primers used in this study**

